# Phylogenetic factorization of compositional data

**DOI:** 10.1101/074112

**Authors:** Alex D Washburne, Justin D Silverman, Jonathan W Leff, Dominic J Bennett, John L. Darcy, Sayan Mukherjee, Noah Fierer, Lawrence A David

**Affiliations:** Nicholas School of the Environment, Duke University, Durham, NC 27708; Cooperative Institute for Research in Environmental Sciences (CIRES), University of Colorado, Boulder, Boulder CO 80309; Program in Computational Biology and Bioinformatics, Duke University, Durham, NC 27708; Medical Scientist Training Program, Duke University, Durham, NC 27708; Center for Genomic and Computational Biology, Duke University, Durham, NC 27708; Department of Molecular Genetics and Microbiology, Duke University, Durham, NC 27708; Department of Earth Science and Engineering, Imperial College London, London UK; Institute of Zoology, Zoological Society of London, London UK; Department of Ecology and Evolutionary Biology, University of Colorado, Boulder, CO 80309; Departments of Statistical Science, Mathematics, and Computer Science, Duke University, Durham, NC 27708

## Abstract

Marker gene sequencing of microbial communities has generated big datasets of microbial relative abundances varying across environmental conditions, sample sites and treatments. These data often come with putative phylogenies, providing unique opportunities to investigate how shared evolutionary history affects microbial abundance patterns. Here, we present a method to identify the phylogenetic factors driving patterns in microbial community composition. We use the method, “phylofactorization”, to re-analyze datasets from human body and soil microbial communities, demonstrating how phylofactorization can be a dimensionality-reducing tool, an ordination-visualization tool, and also mass-produce inferences on the edges in the phylogeny in which meaningful differences arose.

## Background

Microbial communities play important roles in human [6], livestock [16] and plant [3] health, biogeochemical cycles [2, 12], the maintenance of ecosystem productivity, bioremediation, and other ecosystem services. Given the importance of microbial communities and the vast number of uncultured and undescribed microbes associated with animal and plant hosts and in natural and engineered systems, understanding the factors determining microbial community structure and function is major challenge for modern biology.

Marker gene sequencing (e.g. 16S rRNA gene sequencing to assess bacterial and archaeal diversity and 18S markers for Eukaryotic diversity) is now one of the most commonly used approaches for describing microbial communities, quanti-fying the relative abundances of individual microbial taxa, and characterizing how microbial communities change across space, time, or in response to known biotic or abiotic gradients.

Analyzing these data is challenging due to the peculiar noise structure of sequence-count data [30], the inherently compositional nature of the data [15], deciding the taxonomic scale of investigation [7, 8, 31], and the high-dimensionalityof species-rich microbial communities [13]. There is a great need and opportunity to develop tools to more efficiently analyze these datasets and leverage information on the phylogenetic relationships among taxa to better identify which clades are driving differences in microbial community composition across sample categories or measured biotic or abiotic gradients [24]. In this paper, we take on these challenges by developing a means to perform regression of biotic/abiotic gradients on branches in the phylogenetic tree, allowing dimensionality reduction to a series of branches in the phylogeny in a manner consistent with the compositional nature of the data.

Many of these challenges can be resolved by performing regression on clades identified in the phylogeny. Consider a study on the effect of oxazolidinones, which affect gram-positive bacteria, on microbial community composition. Rather than regression of antibiotic treatment on abundance at numerous taxonomic levels, statistical analysis of bacterial communities treated with an oxazolidinone should instantly identify the split between gram-positive and gram-negative bacteria as the most important phylogenetic factor determining response to oxazolidinones. Subsequent factors should then be identified by comparing bacteria within the previously-identified groups: identify clades within gram-positives which may be more resistant or susceptible than the remaining gram-positives. Splitting the phylogeny at each inference and making comparisons within the split groups ensures that subsequent inferences are independent of the gram positive - gram negative split which we have already obtained. All of this analysis must be done consistent with the compositional nature of sequence count data.

Here, we provide a method to analyze phylogenetically-structured compositional data. The algorithm, referred to as “phylofactorization”, iteratively identifies the most important clades driving variation in the data through their associations with independent variables. Clades are chosen based on some metric of the strength or importance of their regressions with meta-data, and subsequent clades are chosen by comparison of sub-clades within the previously-identified bins of phylogenetic groups. Each “factor” identified corresponds to an edge in the phylogeny, and phylofactorization builds on literature from compositional data analysis to construct a set of orthogonal axes corresponding to those edges; the output orthonormal basis allows the projection of sequence-count relative abundances onto these phylogenetic axes for dimensionality reduction, visualization, and standard multivariate statistical analyses. The visualizations and inferences drawn from phylofactorization can be tied back to splits in a given phylogenetic tree and thereby allow researchers to annotate the microbial phylogeny from the results of microbiome datasets.

We show with simulations that phylofactor is able to correctly identify affected clades. We then phylofactor a dataset of human oral and fecal microbiomes to determine the phylogenetic factors driving variation in human body site [4], and a dataset of soil microbes using a multiple regression of pH, carbon concentration and nitrogen concentration [28]. In the human microbiome dataset, we find three splits in the phylogeny that together capture 17.6% of the variation community composition across two body sites. Phylofactorization reveals splits between unclassified OTUs not identifiable by taxonomic grouping, important clades of monophyletic yet para-taxonomic OTUs, and a spectrum of taxonomic scales for binning and analyzing taxonomic units that varies across taxa - all features that would be missed by standard taxonomy-based analysis. In the soil microbiome dataset, we use phylofactor-based dimensionality reduction and ordination-visualization - either using the orthogonal axes corresponding to splits in the phylogeny, or binning OTUs based on their inferred phylogenetic factors - to find that pH drives most of the variation in the dominant clades in the soil dataset, and confirm this finding by dominance analysis on the underlying regressions in phylofactorization, indicating that >%90 of the explained variation in the first three factors is explained by pH. The axes in our ordination-visualization plots correspond to identifiable edges on the phylogeny that have clear biological interpretations and can be used and tested across studies. User-friendly code for implementing, summarizing and visualizing phylofactorization is provided in an R package - ‘phylofactor’.

## Results

We find three main results. First, we find that our algorithm out-performs a standard tool for analyzing compositions of parts related by a tree - what we refer to as the “rooted ILR” transform - and that we can obtain a conservative estimate of the number of phylogenetic factors in simulated datasets a with a known number of affected clades. Second, we phylofactor a dataset of the human oral and fecal microbiomes and find three edges in the phylogeny that account for 17.6% of the variation in microbial communities across these sample sites, edges that are not assigned a unique taxonomic label and are thus invisible to taxonomic-based analyses. Third, we show that phylofactorization can be combined with multiple regression to reveal that pH drives the main phylogenetic patterns of community composition in soil microbiomes, and show that in four factors we split the Acidobacteria three times - including one split that identifies a monophyletic clade of Acidobacteria that consists of alkaliphiles. Finally, using the soil dataset, we demonstrate how phylofactorization yields two complimentary methods for dimensionality reduction and ordination-visualization that tell a simplified story of how the major phylogenetic groups of OTUs change with pH.

### Power Analysis and Conservative Stopping of Phylofactorization

Phylofactorization remedies the structured residuals from the rooted ILR regression on data with fold-changes in abundances within clades. Phylofactorization also remedies the problem of high false-positive rates arising from the nested-dependence and correlated coordinates of the rooted ILR transform, as sequential inferences in phylofactorization are independent. Phylofactorization out-performs the rooted ILR in identifying the correct clades with a given fold-change in abundance (Figs. 1a and 1b), and can be paired with other algorithms assessing residual structure to stop factorization when there is no residual structure and thus accurately identify the number of affected clades (Fig. 1c). Finally, by focusing the inferences on edges instead of nodes in the phylogeny, this algorithm works on trees with polytomies and doesn’t require a forced resolution of polytomies to construct a sequential binary partition of the OTUs. Since edges are the locations of the phylogeny where functional traits arise, the identification of edges that drive variation yields a clear, biological interpretation.

**Figure 1:**
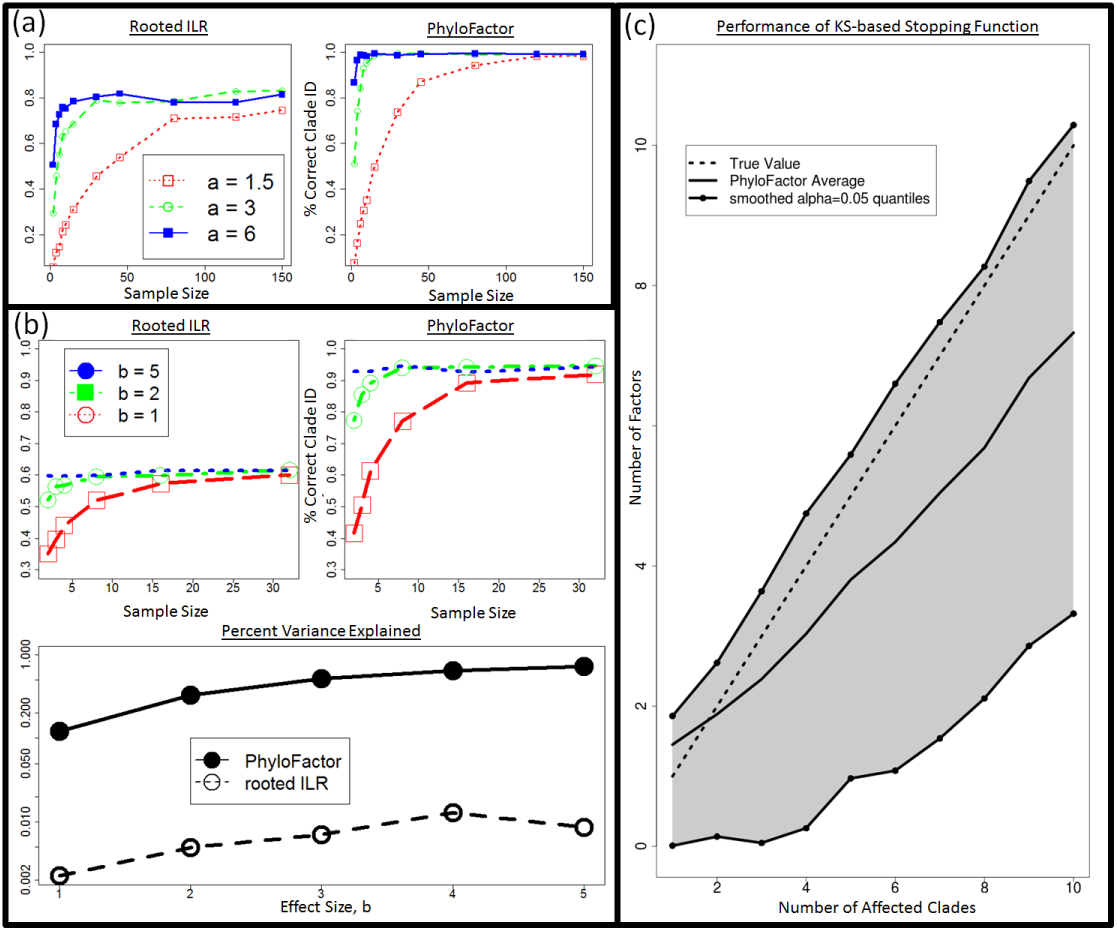
(a) Power Analysis - 1 Clade. The rooted ILR transform that minimizes residual variance when regressed against sample site is less able to identify the correct clade compared to phylofactorization for a variety of effect sizes, a, and sample sizes. (b) Three Significant Clades: When three significant clades are chosen and given a set of effects increasing in intensity with the parameter *b*, choosing the top rooted ILR coordinates under performs phylofactorization in correctly identifying the affected clades. Phylofactorization also explains more variation in the data: across effect sizes, phylofactorization explains 2 orders of magnitude more of the variance in the dataset than the sequential rooted ILR. (c) Stopping Phylofactorization: Plots of the true number of affected clades in simulated datasets against the number of clades identified by the R package ‘phylofactor’. One can terminate phylofactorization when the true number of affected clade is unknown by choosing a stopping function aimed at stopping when there is no evidence of a remaining signal. By stopping the iteration when the distribution of P-values from analyses of variance of regression on candidate ILR basis elements is uniform (specifically, stopping when a KS test against a uniform distribution yields *P >* 0.05), we obtain a conservative estimate of the number of phylogenetic factors in the data.

### Oral-Fecal Microbiome

Phylofactorization of the oral-fecal microbiome dataset, with 290 OTUs and 40 samples, yields three factors that explain 17.6% of the variation in the dataset, factors which correspond to clearly visible blocks in phylogenetic heatmaps of the OTU table (Fig. 1). The factors span a range of taxonomic scales and all of them would be invisible to taxonomic-based analyses. Below, we summarize the factors - the P-values from regression, the taxa split at each factor, the body site associations predicted by generalized linear modeling of the ILR coordinate against body site, and finer detail about the taxonomic identities and known ecology of monophyletic taxa being split. Phylofactorization of these data indicates that a few clades explain a large fraction of the variation in the data, and many more clades can be identified as containing the same intricate detail as the phylogenetic factors presented below. The biology of microbial human-body-site association can focus on these dominant factors - which traits and evolutionary history drive these monophyletic groups’ strong, common association with body sites?

The first factor (*P* = 4.90 *×* 10^*−*30^) split Actinobacteria and Alpha-, Beta-, Gamma-, and Delta-proteobacteria from Epsilonproteobacteria and the rest (Fig. S4). The underlying generalized linear model predicts the Actinobacteria and non-Epsilon-proteobacteria to be 0.4x as abundant as the rest in the gut and 3.7x as abundant as the rest in the tongue. The Actinobacteria identified as more abundant in the tongue include four members of the plaque-associated family Actinomycetaceae, one unclassified species of Cornybacterium) three members of the mouth-associated genus Rothia f20}) and one unclassified species of the vaginal-associated genus Atopobium [9]. With a standard multivariate analysis of the CLR-transformed data, all nine of these Actinobacteria were identified as significantly more abundant in the tongue from regression of the individual OTUs when using either a 1% false-discovery rate or a Bonferonni correction - these monophyletic taxa all individually show a strong preference for the same body site, and their basal branch was identified as our first phylogenetic factor. The remaining Alpha-, Beta-, Gamma- and Delta-proteobacteria grouped with the Actinobacteria consisted of 31 OTUs, and the Epsilonproteobacteira split from the rest were three unclassified species of the genus Campylobacter. The grouping of Actinobacteria with the non-Epsilon Proteobacteria motivates the need for accurate phylogenies in phylofactorization, but also illustrates the promise of identifying clades of interest where the phylogeny is correct and the taxonomy is not.

The second factor (*P* = 1.15 *×* 10^*−*31^) splits 16 Firmicutes of the class Bacilli from the obligately anaerobic Firmicutes class Clostridia and the remaining paraphyletic group containing Epsilonproteobacteria and the rest. The Bacilli are, on average, 0.3x as abundant in the gut as the paraphyletic remaining OTUs and 3.9x as abundant in the tongue. The 16 Bacilli OTUs factored here contain 12 unclassified species of the genus Streptococcus) well known for its association with the mouth [18], one member of the genus Lactococcus, one unclassified species of the mucosal-associated genus Gemella, and two members the family Carnobacteriaceae often associated with fish and meat products [22].

The third factor (*P* = 1.37 *×* 10^*−*28^) separated 15 members of the Bacteroidetes family Prevotellaceae from all other Bacteroidetes and the remaining para-phyletic group of OTUs not split by previous factors. The Prevotellaceae split in the third factor were all of the genus Prevotella) including the species Prevotella melaninogenica and Prevotella nanceiensis found to have abundances 0.3x as abundant in the gut and 4.0x as abundant in the tongue relative to the other taxa from which they were split.

These first three factors capture major blocks visible in the dataset (Fig. 1) can be used as dimensionality reduction tool with a phylogenetic interpretation (Fig. 1). While traditional ordination-visualization tools may capture larger fractions of variation of the data, phylogenetic factorization yields a few variables - ratios of clades - which capture large blocks of variation in the data and can be traced to single edges in the phylogeny corresponding to meaningful splits between taxa, edges where traits likely arose which govern the differential abundances across sample sites and environmental gradients or responses to treatments (Fig. 1b, supplemental Figs. S4-S8).

Using the KS-test stopping criterion, phylofactorization was terminated at 142 factors, each corresponding to a branch in the phylogenetic tree separating two groups of OTUs based on their differential abundances in the tongue and feces. These 142 factors define 143 groups, or what we call ‘bins’, of taxa which remain unsplit by the phylofactorization. The bins vary in size; 112 bins contained only single OTUs, whereas 8 were monophyletic clades and the rest are paraphyletic groups of OTUs, the result of taxa within a monophyletic group being factored, yielding one monophyletic group and one paraphyletic group. Of the 112 single-OTU bins extracted from phylofactorization, 78 were also identified as significant at a false-discovery rate of 1%. Some monophyletic bins included groups of unclassified genera that would not be grouped at the genus level under standard taxonomy-based analyses. For instance, two monophyletic clades of the Firmicutes family Lachnospiraceae were identified as having different preferred body sites, yet both clades were unclassified at the genus level. Taxonomic-based analyses would either omit these unclassified genera, or group them together and make it difficult to observe a signal due to the two sub-groups having different responses to body site.

Performing regression on centered log-ratio (CLR) transformed OTU tables yielded 236 significant OTUs at a false-discovery rate of 1%, and the phylogenetic signal of these OTUs may be difficult to parse out. However, three iterations of phylofactorizaiton yielded the three major splits in the phylogeny, all of which are consistent with known distributions of taxa. Algorithms such as phylosignal [19], which track P-values up the tree, identify clades with common significance, yet not necessarily clades with common signal - it is a common signal, not a common significance, which better indicates a putative trait driving predictable responses in microbes. In the 142 factors above, phylofactor identified numerous clades with common significance yet different signals.

### Soil Microbiome

The soil microbiome dataset was much larger - 3,379 OTUs and 580 samples - and a much smaller fraction of the variation could be explained by phylofactorization. Phylofactorization allows meaningful dimensionality reduction by both factors - plots of the ILR coordinates for the dominant factors - and by bins of taxa that remain un-split at a given level of factorization. Phylofactorization confirmed that the pH of the environment plays a dominant role in the microbial community composition, consistent with previous analyses based on Mantel tests [28]. Dominance analysis of the generalized linear models associated with each factor determined pH to account for approximately 92.87%, 89.78%, and 92.94% of the explained variance in the first, second, and third factor, respectively. C and N were relatively unimportant, and the dominance of pH in the first three factors can be visualized by ordination-visualization plots of the ILR coordinates of the first three factors (Fig. 2a).

**Figure 2:**
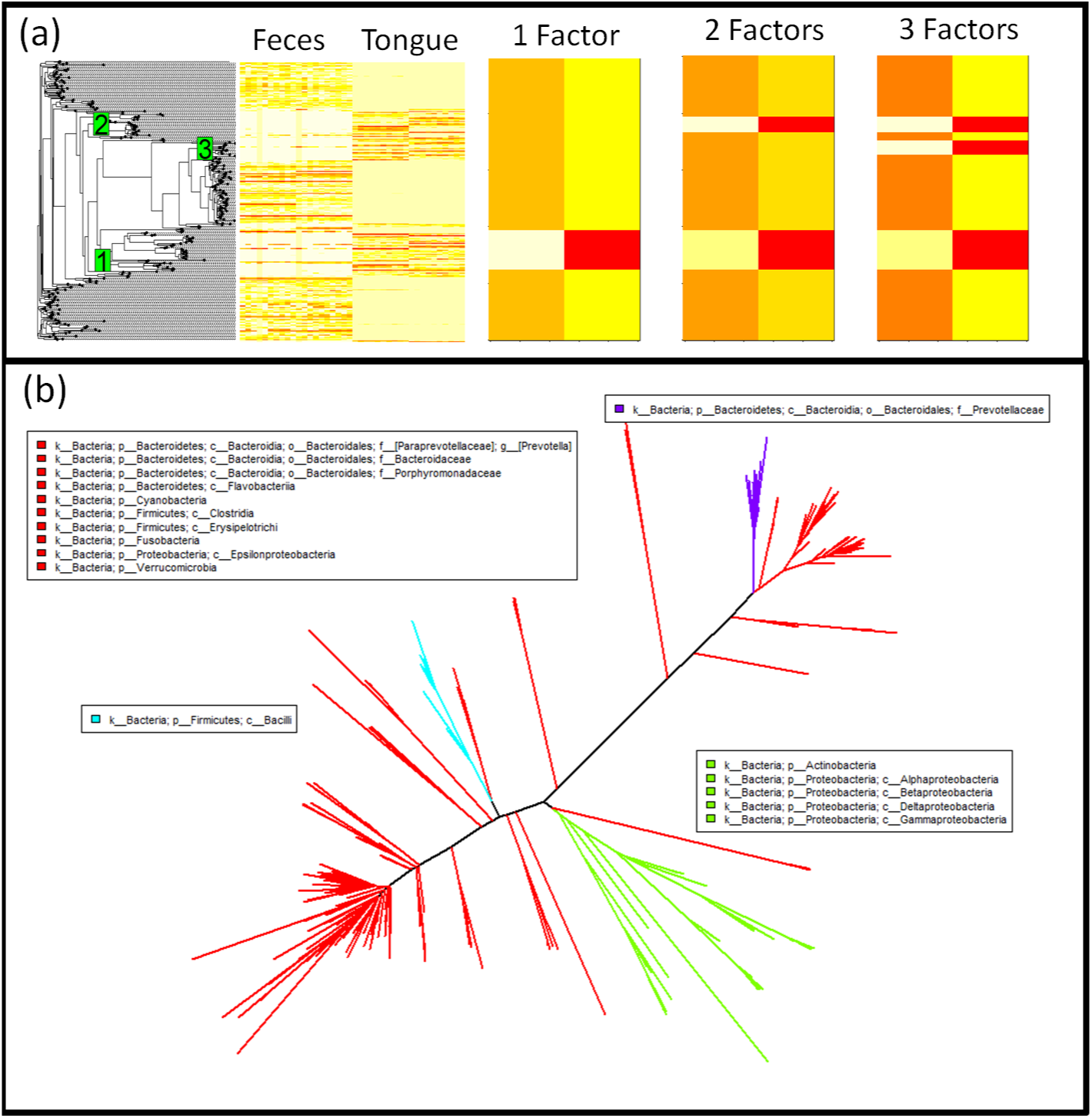
Phylofactorization of human feces/tongue dataset identifies clades differentiating sites. **(a)** Phylogenetic structure is visible as blocks using a phylogenetic heatmap from the R package ‘phytools’ [29]. The first factor separates Actinobacteria and some Proteobacteria from the rest, the second factor separates the class Bacilli from the remaining non-Proteobacteria and non-Actinobacteria, the third factor pulls out the genus Prevotella from Bacteroidetes and indicates that it, unlike many other taxa in Bacteroidetes, is unrepresented in the tongue. Each factor captures a major block of variation in the data, and the orthogonality of the ILR coordinates from each factor allow multiple factors to be combined easily for estimates of community composition. **(b)** These three factors splits the phylogeny into four bins. Three of those bins are monophyletic and the final bin is a “remainder” bin, containing taxa split off by the previous monophyletic bins. The three factors are identifiable edges between nodes that can be mapped to an online database containing those nodes.

The first factor splits a group of 206 OTUs in two classes of Acidobacteria from all other bacteria: class Acidobacteriia and class DA052 are shown to decrease in relative abundance with increasing pH. The second factor split 31 OTUs in the order Actinomycetales (some from the family Thermomonosporaceae and the rest unclassified at the family level) from the remainder of all other bacteria, and these monophyletic Actinomycetales also decrease in relative abundance with increasing pH. The third factor identified another clade within the phylum Acidobacteria to decrease with pH: 115 bacteria from the classes Solibacteres and TM1.

Interestingly, the fourth factor identifies a large collection of 193 OTUs in the remainder of phylum Acidobacteria (i.e. those Acidobacteria not mentioned above in factors 1 and 3) as having relative abundances that increase with pH (dominance analysis: 94.79% of explained variance attributable to pH). Unlike the previous three factors above which were acidophiles, this monophyletic group of Acidobacteria consists of alkaliphiles, which includes the classes Acidobacteria-6, Chloracidobacteria, S053 and three OTUs unclassified at the class level.

The first four factors can be used to define 5 bins of OTUs that we refer to as “binned phylogenetic units” or BPUs: a monophyletic group of Acidobacteria (classes Chloracidobacteria, Acidobacteria-6, and S035), another monophyletic group of Acidobacteria (classes Solibacteres and TM1), a monophyletic group of several families of the order Actinomycetales, a monophyletic group of Acidobacteria (classes Acidobacteriia and DA0522), and a paraphyletic amalagamation of the remaining taxa. Binning the OTUs based on these BPUs tells a simplified story of how pH drives microbial community composition (Fig. 2b).

**Figure 3:**
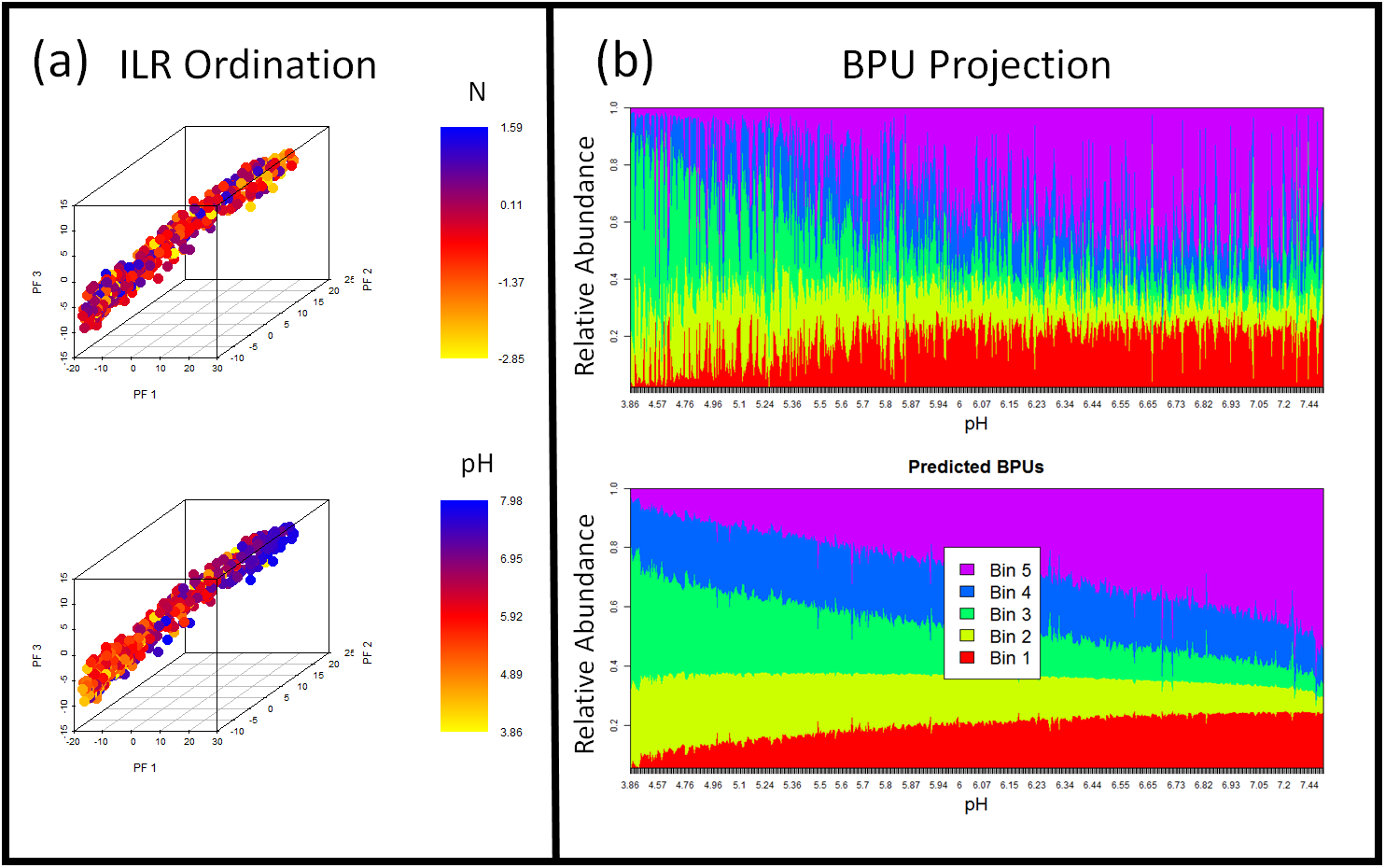
Dimensionality Reduction and Ordination-Visualization of soil microbiome dataset. Phylofactor presents two complementary methods for projecting and visualizing the high-dimensional phylogenetically-structured compositional data. **(a)** The ILR coordinates have asymptotic normality properties and provide biologically informative ordination-visualization plots. Here, we we see that pH is a much better predictor than N of the major phylogenetic factors in Central Park soils. Dominance analysis indicated that pH accounts for approximately 92.87%, 89.78%, and 92.94% of the explained variance in the first, second, and third factor, resepectively, consistent with previous results based on Bray-Curtis distances and Mantel tests showing the dominance of pH in structuring soil microbiomes [28]. **(b)** Every edge separates one group of taxa into two, and those split groups of taxa - what we refer to as bins - can be used to amalgamate taxa and construct a lower-dimensional, compositional dataset of “binned phylogenetic units” (BPUs). Bin 5 is an amalgamation of a monophyletic group of Acidobacteria (classes Chloracidobacteria, Acidobacteria-6, and S035) that increase in relative abundance with pH. Bin 4 is a monophyletic group of Acidobacteria (classes Solibacteres and TMl), Bin 3 is a monophyletic group of several families of the order Actinomycetales, Bin 2 is a monophyletic group of Acidobacteria (classes Acidobacteriia and DA0522), and Bin l is a paraphyletic amalagamation of the remaining taxa.

## Discussion

### Overview

We have introduced a simple and generalizable exploratory data analysis algorithm, phylofactorization, to identify clades driving variation in microbiome datasets. Phylofactorization integrates both the compositional and phylogenetic structure of microbiome datasets and produces outputs that contain biological information: effects of independent variables on edges in the phylogeny, including the tips of the tree traditionally analyzed. The output of phylofactorization contains a sequence of “factors”, or splits in the tree identifying sub-groups of taxa which respond differently to treatment relative to one-another. The splits identified in phylofactorization need not be splits in the Linean taxonomy but can identify strong responses in clades of unclassified taxa. The researcher does not need to choose a taxonomic level at which to perform analysis - those taxonomic levels are output based on whichever clades maximize the objective function, and so researchers will be able to identify multiple taxonomic scales of importance.

Phylofactorization outputs an isometric log-ratio transform of the data with known asymptotic normality properties, coordinates that can be analyzed with standard multivariate methods [25]. The resulting coordinates correspond to particular edges between clearly identifiable nodes in the tree of life, allowing researchers to annotate a given phylogenetic tree with correlations between clades and various environmental meta-data, sample categories, or experimental treatments.

### Future Work

The generality of phylofactorization opens the door to future work employing phylofactorization with other objective functions. As we showed with the human oral/fecal microbiomes, phylofactorization is not restricted to basal clades, but includes the tips as possible clades of interest, but the objective function we used minimized residual variance in the whole community and thereby may prioritize deeply rooted edges or abundant taxa with weaker effects over individual OTUs with stronger effects. Other objective functions could be constructed to meet the needs of the researcher. If researchers are interested in identifying basal lineages, their objective function can weight edges based on distance from the tips. If researchers are interested in identifying putative traits, they may be interested in an objective function weighting edges based on edge length under an assumption that the probability of a trait arising increases with the amount of time elapsed.

Each edge identified in phylofactorization corresponds to two bins of taxa on each side of the edge, and consequently phylofactorization brings in two complementary perspectives for analyzing the data: factor-based analysis and bin-based analysis. Factor-based analysis looks at the each factor as an inference on an edge in the phylogeny, conditioned on the previous inferences already made, and indicating that taxa on one side of an edge respond differently to the independent variable compared to taxa on the other side of the edge. Bin-based analysis, on the other hand, looks at the set of clades resulting from a certain number of factors - what we call a “binned phylogenetic unit” (BPU). These bins will create a lower-dimensional, compositional dataset and can be freed from the underlying ILR coordinates for different analyses on these amalgamated clades. While factor-based analysis provides inferences about the splits in the phylogeny, BPU-based analysis conditions on the factors and bins OTUs based on which factors they contain. BPU-based analysis can inform sequence binning in future research aimed at controlling for previously-identified phylogenetic causes of variation, and combine the effects of multiple up-stream factors for predictions of OTU abundance. See the supplementary text for a more detailed discussion of factor-based and bin-based analyses.

Phylofactorization will benefit from community discussion and further research overcoming general statistical challenges common to greedy algorithms and analysis of phylogenetically-structured compositional data. For instance, the log-ratio transform at the heart of phylofactorization requires researchers deal with zeros in compositional datasets. While there are many methods for dealing with zeros [1, 23, 25], it’s unclear which method is most robust for downstream phylofactorization of sparse OTU tables. Second, phylofactorization as presented here does not allow for multiple regression of ILR basis elements - the set of factors identified after *n* iterations may explain less variation combined than an alternative set of factors that did not maximize the explained variance at each iteration. This limitation may be overcome by running many replicates of a stochastic greedy algorithm and choosing that which maximizes the explained variance after *n* factors. Third, the researcher must choose an objective function which matches her question, and future research can map out which objective functions are appropriate for which questions in microbial ecology. Fourth, like any method performing inference based on phylogenetic structure, phylofactorization assumes an accurate phylogeny. Accurate statistical statements about a researcher’s confidence in phylofactors must incorporate the uncertainty in our constructed phylogeny. Finally, future research can investigate the unique kinds of errors in phylofactorization: in addition to the multiple-hypothesis testing of edges, phylofactorization may propagate errors in the greedy algorithm, and, even when taxa are correctly factored into the appropriate functional bins, the presence of multiple factors in the same region of the tree can lead to uncertainty about the exact edge along which a putative trait arose (see supplement for more discussion on the uncertainty of which edge to annotate).

Incorporating that phylogenetic structure into the analysis of microbiome datasets has been a major challenge [24], and now phylofactorization provides a general framework for rigorous exploration of phylogenetically-structured compositional datasets. The soil dataset analyzed above, for instance, contains 3,379 OTUs and 580 samples, and phylofactorization of the clades affected by pH in the soil dataset yielded not just the three dominant factors used for ordination-visualization, but 2,091 factors in all, each with an intricate phylogenetic story. Many Acidobacteria are acidophiles, but some - Chloracidobacteria, Acidobacteria-6, S035, and some undescribed classes of bacteria factored here - appear to be al-kaliphiles. By incorporating the phylogenetic structure of microbiome datasets, the big data of the modern sequence-count boom just got bigger, and future research will need to consider how to organize, analyze and visualize the large amounts of phylogenetic detail that can now be obtained from the analysis of microbiome datasets.

## Conclusions

Phylofactorization is a robust tool for analyzing marker gene sequence-count datasets for phylogenetic patterns underlying microbial community responses to independent variables. Phylofactorization accounts for the compositional nature of the data and the underlying phylogeny and produces inferences that are independent and more powerful than application of the ILR transform to the rooted phylogeny. The R package ‘phylofactor’ has built-in parallelization that can be used to analyze large microbiome datasets, and allows generalized linear modeling to identify clades which respond to treatments or multiple environmental gradients.

Phylofactorization can connect the pipeline of microbiome studies to focused studies of microbial physiology. As researchers identify lineages with putative functional ecological responses, taxa within those lineages - even if they are not the same OTUs - can be cultivated and their genomes screened to uncover the physiological mechanisms underlying the lineages’ shared response.

Phylofactorization improves the pipeline for analyzing microbiome datasets by allowing researchers to objectively determine the appropriate phylogenetic scales for analyzing microbiome datasets - a family here, an unclassified split there - instead of performing multiple comparisons at each taxonomic level. Instead of principle components analysis or principle coordinates analysis, phylofactorization can be used as for exploratory data analysis and dimensionality reduction tool in which the “components" are identifiable clades in the tree of life, a far more intuitive and informative component for biological variation than multispecies loadings.

Phylofactorization can allow researchers to annotate online databases of the microbial tree of life, permitting predictions about the physiology of unclassified and uncharacterized life forms based on previous phylogenetic inferences in sequence-count data. By allowing researchers to make inferences on the same tree and potentially annotate an online tree of life, phylofactorization may bring on a new era of characterizing high-throughput phylogenetic annotations, filling in the gaps the microbial tree of life.

An R package for phylofactorization with user-friendly parallelization is now available online at https://github.com/reptalex/phylofactor.

## Methods

### Phylogenetically-Structured compositional data

Microbiome datasets are “phylogenetically-structured compositional data”, compositions of parts linked together by a phylogeny for which only inferences on relative abundances can be drawn. The phylogeny is the scaffolding for the evolution of vertically-transmitted traits, and vertically-transmitted traits may underlie an organism’s functional ecology and response to perturbations or environmental gradients. Performing inference on the edges in a phylogeny driving variation in the data can be useful for identifying clades with putative traits causing related taxa to respond similarly to treatments, but such inferences must account for the compositional nature of the sequence-count data.

A standard analysis of microbiome datasets uses only the distal edges of the tree - the OTUs - and a few edges within the tree separating Linean taxonomic groups. However, a phylogeny of *D* taxa and no polytomies is composed of 2*D −* 3 edges, each connecting two disjoint sets of taxa in the tree with no guarantee that splits in Linean taxonomy corresponds to phylogenetic splits driving variation in our dataset. Thus, instead of analyzing just the tips and a series of Linean splits in the tree, a more robust analysis of phylogenetically-structured compositional data should analyze all of the edges in the tree. To do that, we draw on the isometric log-ratio transform from compositional data analysis, which has been used to search for a taxonomic signature of obesity in the human gut flora [14] and incorporated into packages for downstream principal components analysis [21]. However, to the best of our knowledge, the previous literature using the isometric log-ratio transform in microbiome datasets has used random or standard sequential binary partitions, and not explicitly incorporated the phylogeny as their sequential binary partition.

### The Isometric Log-Ratio Transform of a rooted phylogeny

The isometric log-ratio (ILR) transform was developed as a way to transform compositional data from the simplex into real space where standard statistical tools can be applied [11, 10]. A sequential binary partition is used to construct a new set of coordinates, and the phylogeny is a natural choice for the sequential binary partition in microbiome datasets. Instead of analyzing relative abun-dances, *y*_*i*_, of *D* different OTUs, the ILR transform produces *D −* 1 coordinates, 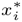(called “balances”). Each balance corresponds to a single internal node of the tree and represents the averaged difference in relative abundance between the taxa in the two sister clades descending from that node (the difference being appropriately measured as a log-ratio due to the compositional nature of the data; see SI for more detailed description of the ILR transform). For an arbitrary node indicating the split of a group, *R* with *r* elements from the group, *S* with *s* elements, the ILR balance can be written as

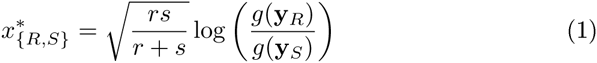

where *g*(**y**_*R*_) is the geometric mean of all *y*_*i*_ for *i ∈ R*.

We refer to the ILR transform corresponding to a rooted phylogeny as the “rooted ILR”. The rooted ILR creates a set of ILR coordinates, 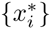, where each coordinate corresponds to the “balance” between sister clades at each split in the phylogenetic tree. The balances in a rooted ILR transform in equation (1) can be intuited as the average difference between taxa in two groups, and splits in the tree which meaningfully differentiate taxa will be those splits in which the average difference between taxa in two groups changes predictably with an independent variable. Inferences on ILR coordinates, then, map to inferences on lineages in the phylogenetic tree.

The rooted ILR coordinates provide a natural way to analyze microbiota data as they measure the difference in the relative abundances of sister clades and may be useful in identifying effects contained within clades such as zero-sum competition of close relatives or the substitution of one relative for another across environments. However, if we desire to link the effect of an external covariate (e.g. antibiotics vs. no antibiotic treatment) to clades within the phylogeny, the best comparison may not be between sister clades, but instead between all other clades, controlling for any other phylogenetic splits or factors we may know of (e.g. we may compare a lineage within gram-positives with all other gram-positives, once we’ve identified the gram-positive vs. gram-negative split as an important factor for antibiotic susceptibility). We refer to this unrooted approach as ‘phylofactorization’.

For the task of linking an external covariate to individual clades in the phylogeny, we examine three features of the rooted ILR that can be improved on by phylofactorization by considering a treatment that decreases the abundance of one and only one clade, *B*, whose closest relative is clade *A*. Regression on the rooted ILR coordinates may identify the balance 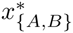 corresponding to the most recent common ancestor of clades *A* and *B* as having that strongest response to the treatment, but regression on this coordinate will suggest that clade *B* decreases relative to *A*, leading to structured residuals in the original dataset due to an inability to account for the increase in clade *B* relative to the rest of the OTUs in the data (Fig. 4a). Second, all partitions between affected clade and the root will be affected. If each balance is tested independently, the rooted ILR may identify many clades that are affected by antibiotics; the correlations between coordinates can yield a high false-positive rate if just one clade is affected (Fig. 4b). Finally, the ILR transformation does not work with polytomies common in real, unresolved phylogenies. Any polytomy will produce a split in the phylogeny between three or more taxa, and there is no general way to describe the balance of relative abundances of three or more parts using only one coordinate.

**Figure 4:**
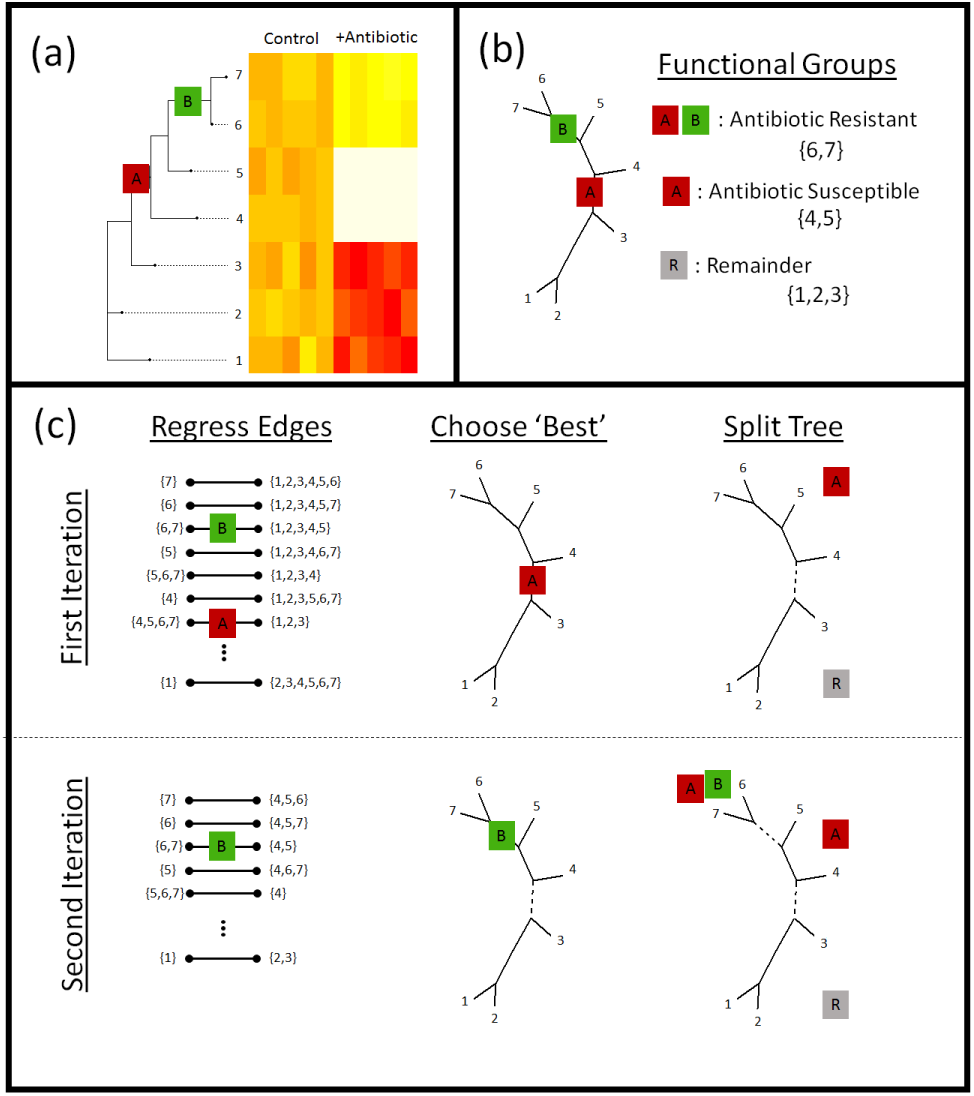
Phylofactorization: **(a)** Phylofactorization changes variables from tips of the phylogeny (OTUs used in analysis of microbiome datasets) to edges of the phylogeny with the largest predictable differences between taxa on each side of the edge. To illustrate this method, we consider the treatment of a bacterial community with an oxazolidinone. Oxazolidinones target gram-positive bacteria and will likely lead to a decrease in the relative abundances of gram-positive bacteria (antibiotic susceptible clade, A, having the antibiotic target). Among the antibiotic susceptible bacteria, phylofactor can identify monophyletic clades that are resistant relative to other antibiotic-susceptible bacteria due to a vertically-transmitted trait (B) such as the loss of the antibiotic target or enzymes that break down the antibiotic. **(b)** The two phylogenetic factors produce three meaningful bins of taxa - those susceptible to antibiotics (A), those within the susceptible clade that are resistant to antibiotics (A+B), and a potentially paraphyletic remainder. **(c)** Phylofactorization is a greedy algorithm to extract the edges which capture the most predictable differences in the response of relative abundances among taxa on the two sides of each edge. (c, top row) For the first iteration, all edges are considered - an ILR coordinate is created for each edge using equation and the ILR coordinate is regressed against the independent variable. The edge which maximizes the objective function is chosen. Depicted above, the first factor corresponds to the edge separating antibiotic susceptible bacteria from the rest. Then, the tree is split - all subsequent comparisons along edges will be contained within the sub-trees. The conceptual justification for limiting comparisons within sub-trees is to prevent over-lapping comparisons: once we identify the antibiotic susceptible clade, we want to look at which taxa within that clade behave differently from other taxa within that clade. (c, bottom row) For the second iteration, the remaining edges are considered, ILR coordinates within sub-trees are constructed. The edge maximizing the objective function is selected and the tree is split at that edge. For more details, see the section “PhyloFactor" in the supplemental info.

Nonetheless, the simplicity and theoretical foundations underlying the ILR, and the instant appeal of applying it to the sequential-binary partition of the phylogeny, motivate the rooted ILR as a simple tool for analysis of the phylogenetic structure in compositional data. For that reason, we use the rooted ILR as a baseline for comparison of our more complicated method of phylogenetic factorization.

### Phylofactorization

The shortcomings of the rooted ILR can be remedied by modifying the ILR transform to apply not to the nodes or splits in a phylogeny, but to the edges in an unrooted phylogeny. While ILR coordinates of nodes allow a comparison of sister clades, ILR coordinates along edges allow comparison of taxa with putative traits that arose along the edge against all taxa without those putative traits. Traits arise along edges of the phylogeny and so, for annotation of online trees of life, effects in a clade are best mapped to a chain of edges in the phylogeny. However, the ILR transform requires a sequential binary partition, and the edges don’t immediately provide a clear candidate for a sequential binary partition. In what we refer to as “phylofactorization”, one can iteratively construct a sequential binary partition from the unrooted phylogeny by using a greedy algorithm by sequentially choosing edges which maximize a researcher’s objective function. Phylofactorization consists of 3 steps (Box 1): (1) Consider the set of possible primary ILR basis elements corresponding to a partition along any edge in the tree (including the tips). (2) Choose the edge whose corresponding ILR basis element maximizes some objective function - such as the test-statistic from regression or the percent of variation explained in the original dataset - and the groups of taxa split by that edge form the first partition. (3) Repeat steps 1 and 2, constructing subsequent ILR basis elements corresponding to remaining edges in the phylogeny and made orthogonal to all previous partitions by limiting the comparisons to taxa within the groups of taxa un-split by previous partitions.

**Figure 5:**
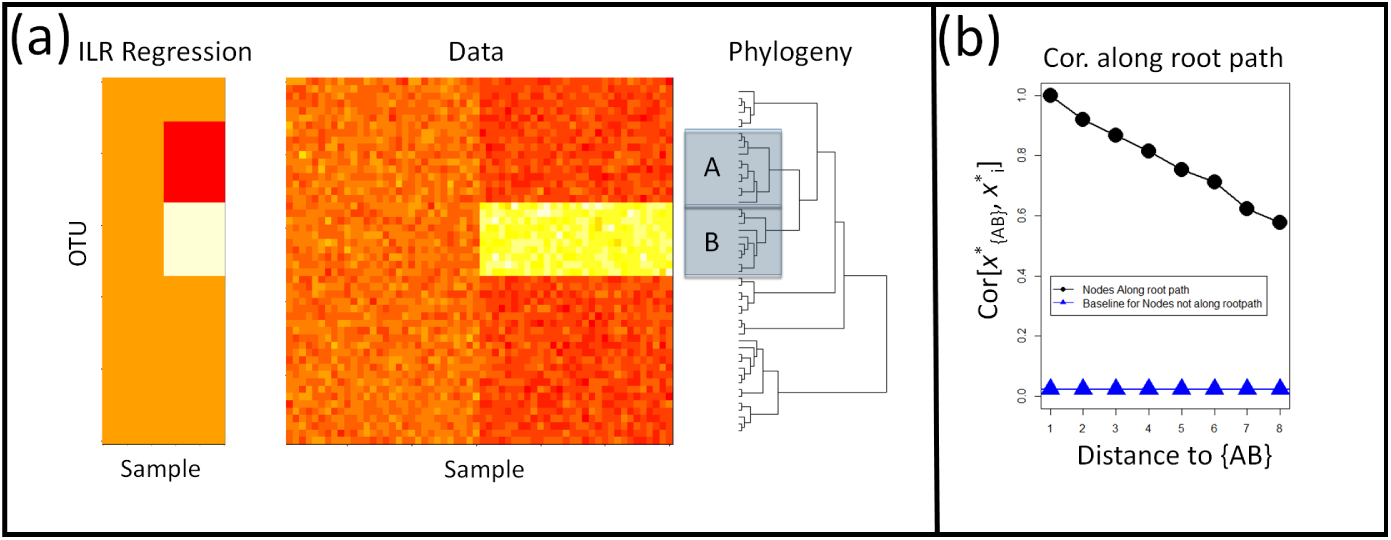
Shortcomings of Rooted ILR. **(a)** The isometric log-ratio transform corresponding to a phylogeny rooted at the common ancestor is inaccurate for geometric changes within clades. Here, absolute abundances of 50 taxa in 30 samples per site were simulated across two sites. An affected clade, *B*, is up-represented in the second site. Regression on the rooted ILR coordinates, 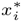, against the sample site indicated that the partition separating clade *A, B*, referred to as 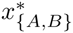, had the highest test-statistic, but the rooted ILR predicts fold-changes in *B* relative to *A*, not fold changes in *B* relative to the rest of the taxa. (b) Consequently, when one clade increase in abundance while the rest remain unaffected, partitions between the affected clade and the root will also have a signal leading to a correlation in the coordinates along the path from *B* to the root. The correlation plotted here is the absolute value of the correlation coefficient, and the baseline correlation was estimated as the average absolute value of the correlation coefficient between ILR coordinates not along the root-path of the affected clade.

Explicitly, the first iteration of phylofactorization considers a set of candidate ILR coordinates, 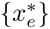 corresponding to the two groups of taxa split by each edge, *e*. Then, regression is performed on each of the ILR coordinates, 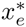 ~ *f(X)* for an appropriate function, *f* and a set of independent variables, *X*. The edge, 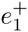, which maximizes the objective function is chosen as the first phylogenetic factor. In this paper, our objective function is the difference between the null deviance of the ILR coordinate and the deviance of the generalized linear model explaining that ILR coordinate as a function of the independent variables. We use this objective function as a measure of the amount of variance explained by regression on each edge because the total variance in a compositional dataset is constant and equal to the sum of the variances of all ILR coordinates corresponding to any sequential binary partition. Consequently, at each iteration there is a fixed amount of the total variance remaining in the dataset, and so at the candidate ILR coordinate which captures the greatest fraction of the total variance in the dataset is the one with the greatest amount of variance explained by the regression. After identifying 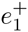, we cut the tree in two sub-trees along the edge, 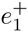.

For the second iteration, another set of candidate ILR coordinates is constructed such that their underlying balancing elements are orthogonal to the first ILR coordinate. Orthogonality is ensured by constructing ILR coordinates contrasting the abundances of taxa along each edge, restricting the contrast to all taxa within the sub-tree in which the edge is found. A new edge, 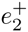, which maximizes the objective function is chosen as the second factor, the sub-tree containing this edge is cut along this edge to produce two sub-trees, and the process is repeated until a desired number of factors is reached or until a stopping criterion is met. More details on the algorithm, along with a discussion on objective functions, is contained in the SI.

While one could use other methods of amalgamating abundances along edges, the conceptual importance of using the ILR transform is twofold: the ILR transform has proven asymptotic normality properties for compositional data to allow the application of standard multivariate methods [11], and the ILR transform serves as a measure of contrast between two groups. The log-ratio used in phylofactor is an averaged ratio of abundances of taxa on two sides of an edge (see supplement for more detail), thus phylofactorization searches the tree for the edge which has the most predictable difference between taxa on each side of the edge, or, put differently, the edge which best differentiates taxa on each side. Thus, each edge that differentiates taxa and their responses to independent variables is considered a phylogenetic “factor" driving variation in the data.

The output of phylofactorization is a set of orthogonal, sequentially “less important” ILR basis elements, their predicted balances, and all other information obtained from regression. After the first iteration of phylofactorization, we are left with an ILR basis element corresponding to the edge which maximized our objective function and split the dataset into two disjoint sub-trees, or sets of OTUs that we henceforth refer to as “bins”, and we have an estimated ILR balancing element, 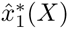, where *X* is our set of independent variables. Subsequent factors will split the bins from previous steps, and after *n* iterations one has *n* factors that can be mapped to the phylogeny, *n* + 1 bins for binning taxa based on their phylogenetic factors, *n* estimates of ILR balancing elements, and an orthonormal ILR basis that can be used to project the data onto a lower dimensional space. The sequential splitting of bins in phylofactorization ensures sequentially independent inferences - having already identified group *B* as hyper-abundant relative to group *A* in the example illustrated in Fig. 4, downstream factors must analyze sub-compositions entirely within *B* and within *A*.

### Computational Tools

Phylofactorization was done using the R package “phylofactor" available at https://github.com/reptalex/phylofactor. The R package contains detailed help files that demo the use of the package, and the exact code used in analyses and visualization in this paper are available in the supplementary materials. The rooted ILR transform was performed as described in [10] where the sequential binary partition was the rooted phylogeny.

### Power Analysis of Rooted ILR and Phylofactorization

To compare the ability of phylofactorization and the rooted ILR to identify clades of OTUs with shared associations with independent variables, we simulated random communities of *D* = 50 OTUs and *p* = 40 samples by simulating random absolute abundances, *N*_*i,j*_, such that log *N*_*i,j*_ were i.i.d Gaussian random variables with mean *µ* = 8 and standard deviation *σ* = 0.5. The OTUs were connected by a random tree (the tree remained constant across all simulations), and then either 1 or 3 clades were randomly chosen to have associations with a binary “environment" independent variable with *p* = 20 samples for each of its two values to represent an equal sampling of microbial communities across two environments.

For simulations with one significant clade, the abundances of all the OTUs within that clade increased by a factor a in the second environment where a *∈* {1.5, 3, 6*}*. For simulations with three significant clades, the three clades were drawn at random and randomly assigned a fold-change from the set *{π*^*b*^, 0.5^*b*^, exp(*−b*)} in a randomly chosen environment where *b* ∈ {1, 2, 5}. For each fold-change, 500 replicates were run to compare the power of the rooted ILR and phylofac-torization in correctly identifying the affected clades.

Regression of rooted ILR coordinates was performed and the coordinates were ranked by the difference between their null deviance and the model deviance. The ability of a rooted ILR coordinate to identify the correct 1 clade or 3 clades was measured by the percent of its top 1 or 3 ILR coordinates, respectively, which corresponded to the node on the tree from which the affected clade(s) originated. The ability of phylofactor to identify the correct 1 clade or 3 clades was measured by the percent of the factors that correctly split an affected clade from the rest (e.g. the percent of factors corresponding to edges along which a trait arose).

For the 3 clade simulations, we also compared the amount of variance explained by 3 factors in phylofactorization with the amount of variance explained by the top 3 ILR coordinates in the rooted ILR. The amount of variance explained was measured as the difference in the null deviance and the model deviance, summed across all three factors or the top 3 ILR coordinates.

### KS-based Stopping Function for PhyloFactor

While a researcher can iterate through phylofactorization until a full basis of *D −* 1 ILR coordinates is constructed, there is value in stopping the iteration when all of the clades have been identified or at a conservative underestimate of the true number of phylogenetic factors. We implemented a stopping function based on a Kolmogorov-Smirnov (KS) test of the distribution of P-values from analyses of variance of the regressions on candidate ILR coordinates. If there is no phylogenetic signal, we anticipate the true distribution of P-values to be uniform (albeit with some dependence among the P-values due to overlap in the OTUs used in the ILR coordinates). Thus, we tested the ability of phylofactor to correctly identify the number of clades if phylofactorization is stopped when a KS test of the P-values produces its own P-value *P*_*KS*_ > 0.05.

We simulated 300 replicate communities with *M* clades for each *M ∈ {*1*, …,* 10}. For simulations with *M* clades, *D* = 50 and *p* = 40 communities were simulated as above and fold changes, *c*, were drawn as log-normal random variables where log(*c*_*k*_) were i.i.d Gaussian random variables with *µ* = 0 and *σ* = 3 for *k* = 1, …, *M*. The number of clades identified by phylofactor for a given true number of clades, *K*_*M,r*_, was tallied for *r* = 1, …, 300. We calculate the mean 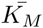 across all replicates and, for visualization purposes, interpolate the *α* = 0.025 and *α* = 0.975 quantiles by finding the best fit of a logistic function to the cumulative distribution of 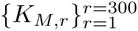 for each *M*.

### Analysis of Fecal/Oral microbiome data

16S amplicon sequencing data from Caporaso et al. (2011) [4] were downloaded from the MG-RAST database (http://metagenomics.anl.gov/) along with associated metadata. QIIME [5] was used to trim primers from these data, and to cluster OTUs with the Greengenes reference database (May 2013 version; http://greengenes.lbl.gov). Longer sequence lengths in the greengenes database (~1400 BP) compared to the original Illumina sequences (~123 BP) allows more informative base pairs for phylogenetic tree construction. We used the phylogenetic tree that is included with the greengenes database for all analyses. The resulting OTU table was rarefied to 6000 sequences per sample.

10 time points were randomly drawn from each of the male tongue, female tongue, male feces and female feces datasets, giving a total of n-20 samples at each site. Taxa present in fewer than 30 of the 40 samples were discarded, and phylofactorization was done by adding pseudo-counts of 0.65 to all 0 entries in the dataset [1], converting counts in each sample to relative abundances, and then regressing the ILR coordinates against body site. The complete R script is available in the file “Data Analysis pipeline of the FT microbiome”.

Complete phylofactorization of this dataset was performed by stopping the algorithm when a KS-test on the uniformity of P-values from analyses of variance of regression on candidate ILR-coordinates yielded *P*_*KS*_ > 0.05. These results were compared with a standard, multiple hypothesis-testing analysis of CLR-transformed data. The summary of the taxonomic detail at the first three factors is provided in the results section, and a full list of the taxa factored at each step is available in the supplement and can be further explored using the R pipeline provided.

### Analysis of Soil microbiome data

The soil microbiome dataset from [28] was included to illustrate the ability of phylofactor to work on bigger microbiome datasets with continuous independent variables and multiple regression. Details on sample collection, sequencing, meta-data measurements and OTU clustering are available in [28]. The phylogeny was constructed by aligning representative sequences using SINA [27], trimming bases that represented gaps in *≥*20% of sequences, and using fasttree

[26].

The complete dataset contained 123,851 OTUs and 580 samples. Data were filtered to include all OTUs with on average 2 or more sequences counted across all samples, shrinking the dataset to D=3,379 OTUs. The data were further trimmed to include only those samples with available pH, C and N meta-data, reducing the sample size to n-551.

Phylofactorization was done by adding pseudo-counts of 0.65 to all 0 entries in the dataset [1], converting counts in each sample to relative abundances, and performing multiple regression of pH, C and N on ILR coordinates. The first three factors are used for ordination-visualization. To determine the relative importance of each abiotic variable in driving phylogenetic patterns of microbial community composition, we used the lmg method from the R package ‘relaimpo’ [17] which averages the sequential sums of squares over all orderings of regressors to obtain a measure of relative importance of each regressor in the multivariate model.

## Acknowledgments

ADW would like to acknowledge L. Ma for his feedback and help incorporating this method into the statistical literature. JS was supported in part by the Duke University Medical Scientist Training Program. This paper is published by support from and in loving memory of D. Nemergut.

## Declarations

**Competing Interests:** The authors have no competing interests in relation to this work.

### Availability of Data and Materials

The data were obtained from previous studies and are available online through the original studies. The R package ‘phylofactor’ is available at https://github.com/reptalex/phylofactor and all other R files used in the analysis and visualization are available online.

